# Trapped pore waters in the open proton channel H_V_1

**DOI:** 10.1101/2022.05.09.491141

**Authors:** Danila Boytsov, Stefania Brescia, Gustavo Chaves, Sabina Koefler, Christof Hannesschlaeger, Christine Siligan, Nikolaus Gössweiner-Mohr, Boris Musset, Peter Pohl

## Abstract

The voltage-gated proton channel, H_V_1, is crucial for innate immune responses. According to alternative hypotheses, protons either hop on top of an uninterrupted water wire or bypass titratable amino acids, interrupting the water wire halfway across the membrane. To distinguish between both hypotheses, we estimate the water mobility for the putative case of an uninterrupted wire. The predicted single-channel water permeability 3×10^−12^cm^3^s^−1^ reflects the permeability-governing number of hydrogen bonds between water molecules in single-file configuration and pore residues. However, the measured unitary water permeability does not confirm the prediction, i.e., it is negligible. Osmotic deflation of reconstituted lipid vesicles reveals trapped water inside the H_V_1 wild-type channel and D174A mutant open at 0 mV. The conductance of 1400 H^+^ s^−1^ per wild-type channel agrees with the calculated diffusion limit for a ~2 Å capture radius for protons. Removal of a charged amino acid (D174) at the pore mouth decreases H^+^ conductance, conceivably by reducing the capture radius. At least one intervening amino acid contributes to H^+^ conductance while blocking water flow.

## Introduction

Many different cell types express the voltage-gated proton channel H_V_1 (*1*). Among other roles, it is involved in innate immunity (*2*) - its primary function being the acid extrusion from cells. Transmembrane voltage and pH gradient contribute to H_V_1 gating (*3*). Its structure is homologous to classical voltage-gated channels’ voltage-sensing domain (VSD) (*4*). Experimentally, exclusively closed structures are available – a crystal structure of a chimera between mouse H_V_1 and *Ciona intestinalis* voltage-sensing phosphatase (*5*) and an NMR structure of the human H_V_1 (*6*). Also, electron paramagnetic resonance (EPR) spectroscopy data decipher the resting structure of H_V_1 (*7*). Recently, an AlphaFold structure has been added (Fig. 1). The open channel’s structure has solely been computed as homology models (*8–14*). Therefore, the molecular mechanism of proton transfer is still debated (*15, 16*).

**Fig. 1.**
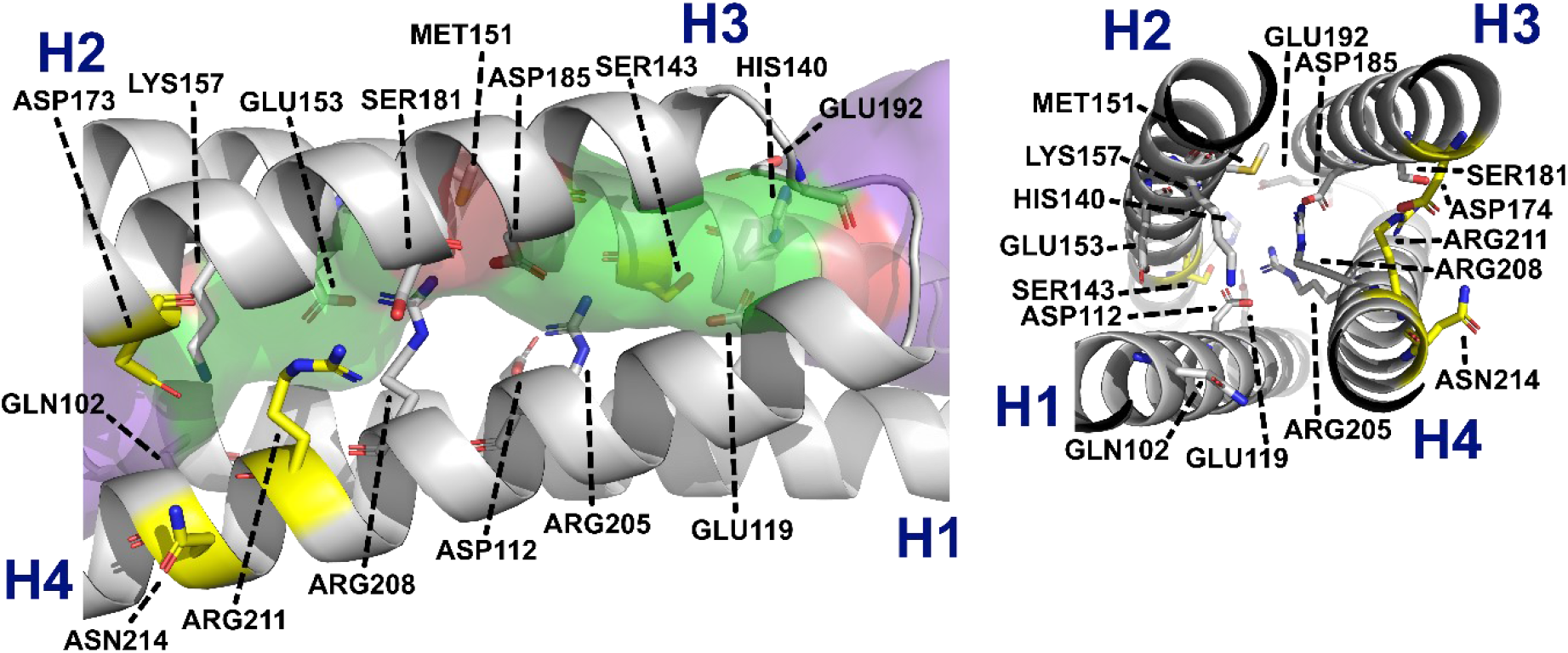
Structure of the H_V_1 channel as predicted by AlphaFold (DOI: 10.1038/s41586-021-03819-2 for AlphaFold /// database entry Q96D96. (A) 12 potential H-bond donors and acceptors guide water through the H_V_1 pore from the cytoplasm (left) to the periplasm (right). We generated the pore surface with the program HOLE (27): Green areas represent pore radii suitable for a single water molecule only, red areas are too narrow to let water molecules pass, and purple regions can accommodate more than one water molecule per cross-section. D112 fixes R205 in a conformation that further constricts the channel. These 12 amino acids and 4 additional residues (colored in yellow: S143, D173, R211, N214) may form hydrogen bonds with pore waters in the single file region. The side chains of the additional amino acids face towards the channel surface in open/closed models as shown in Fig. S1. (B) View along the channel (cyto > peri).

The first hypothesis **α** of proton permeation stipulates that charged residues interrupt the H_V_1 spanning water wire, i.e., water molecules are trapped inside the H_V_1 channel (*10*). In contrast to water, protons can still traverse the channel by hopping from one acidic residue to the next through one or more bridging water molecules (*11*) or, vice versa, by hopping along water wires linked by titratable residues (*17, 18*).

According to the second hypothesis **β**, protons move through the open H_V_1 by hopping over an uninterrupted file of water molecules (*19*), i.e., the same way they move through the peptide channel gramicidin A, gA (*20*), via Grotthuss’ mechanism (*21*). Molecular dynamics simulations based on the K_v_1.2–K_v_2.1 paddle-chimera VSD support the view by showing an uninterrupted water wire that extends through the membrane (*9*).

Thus far, only indirect experimental evidence is available to distinguish between hypotheses **α** and **β**. Comparing deuterium ion and proton currents through the plasma membrane of rat alveolar epithelial cells, DeCoursey & Cherny (*22*) found an isotope effect exceeding that for hydrogen bond cleavage in bulk water. It suggested the involvement of an amino acid side chain in proton conduction (*22*). Alternatively, altered properties of confined water could have been responsible for the higher isotope effect.

We now aim to provide direct experimental evidence for the involvement of one or several amino acid side chains in proton transport. Therefore, we make use of the available insight into channel geometry. Significantly, the intraluminal water molecules must arrange in a single file, i.e., the H_V_1 pore is so narrow that the molecules cannot overtake each other (Fig. 1). The rate at which water crosses the single-file region can be predicted from the number, *N*_H_, of hydrogen bonds that the water molecules may form with pore-lining residues (*23, 24*). Rate’s dependency on *N*_H_ has been established by analyzing other physiologically essential channels which harbor single water files: gramicidin–A (gA), aquaporins, and potassium-selective channels (*25*). They all facilitate water transport (*26*): the larger *N*_H_, the smaller the unitary water permeability, *p*_f_. According to the AlphaFold structure, H_V_1 offers 16 residues, which may form hydrogen bonds with pore waters (Fig. 1). A homology model optimized by microsecond long molecular dynamic simulations (*13*) suggests a smaller number. Only six hydrogen bond-forming residues localize into the single file region of the open conformation (Fig. S1). *N*_H_ = 6 is true also for the bacterial glycerol facilitator GlpF (*23*). GlpF is one of the most efficient water channels known, i.e., it is the single-file water channel with the highest unitary water permeability, *p*_f_ (*26*). However, analyzing the open conformation of the homology model by the HOLE program (*27*) reveals a constriction side too narrow for water molecules to pass (Fig. S1). Surprisingly, the closed conformation of the homology model appears wide enough for water passage (Fig. S1). Yet, the prediction of AlphaFold is different. AlphaFold stipulates that H_V_1 does not offer a water pathway in the closed state (Fig. 1). Below, we show negligible *p*_f_ in H_V_1’s open and closed states, thereby confirming hypotheses **α** and supporting the AlphaFold prediction.

## Results

We purified (Fig. S2) and reconstituted H_V_1 channels to perform water flux measurements. The experimental design requires the channel to be in its open configuration at zero transmembrane potential. Previous experiments suggest that the D174A mutant meets the requirement (*19*). We confirmed the observation by performing patch-clamp studies in whole-cell configuration and comparing D174A to wild-type H_V_1 (Fig. 2).

**Fig. 2.**
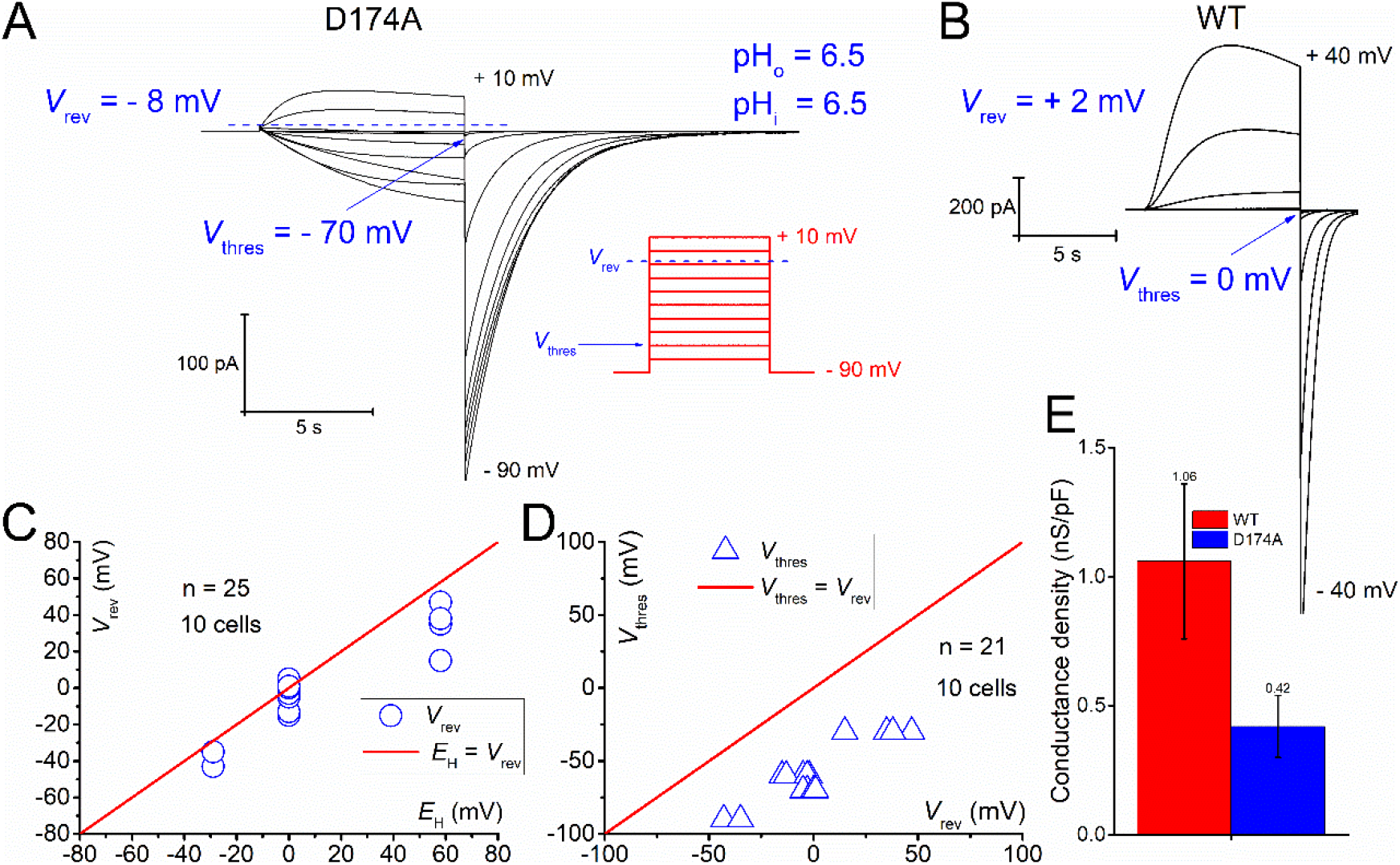
Typical whole-cell patch-clamp measurement of the human H_V_1 D174A mutant in tSA201 cells. **(A)** Family of proton currents due to depolarizing pulses from – 90 mV to + 10 mV, in 10 mV increments, at pH_o_ 6.5 and pH_i_ 6.5. Pronounced inward H^+^ currents activate negatively (here – 70 mV) to reversal potential (here – 8 mV), indicating a high open probability of the D174A mutant at 0 mV. **(B)** Wild type H_V_1 activates close to the reversal potential and depicts a low open probability at 0 mV. **(C)** Comparison of calculated Nernst potential for protons (*E*_H_) and measured reversal potential (*V*_rev_) for the D174A mutant. The red line indicates perfect proton selectivity. **(D)** Comparison of the threshold of activation (*V*_thres_) and measured reversal potential (*V*_rev_) of the D174A mutant, indicating a high open probability without application of external voltage. **(E)** Comparison of the maximal conductance density of cells expressing wild-type H_V_1 and the D174A mutant.

### Whole-cell patch-clamp

We overexpressed wild-type H_V_1 and D174A mutant in tsa201 cells. Using our previous protocol under symmetrical pH conditions (*28, 29*), we detected threshold potentials of activation, *V*_thres_, of – 70 mV for the mutant and 0 mV for the wild type (Fig. 2). At the same time, the D174A mutant retains nearly perfect proton selectivity, i.e., the measured reversal potential was close to the Nernst potential for protons (Fig. 2C). Repeated experiments at different pHs showed that the threshold potential of the mutant is always more negative than the reversal potential for protons.

Consequently, the mutant channel is nearly fully open (Fig. 2D), readily seen when the membrane potential is 0 mV and external voltage is absent. The high open probability of the D174 mutant under symmetrical pH conditions is readily seen in the tail current amplitude reaching a quasi-saturation (Fig. 2A). The resulting outward currents have a higher amplitude in the wild-type (Fig. 2A+B). Interestingly, the conductance density mediated by the expression of the mutant was 2.5 times smaller than the wild type, although we transfected the same amount of plasmid DNA (Fig. 2E). Our observation suggests a reduced flux through the mutant if we assume that protein abundance in the plasma membrane is independent of the mutation.

### Vesicular proton uptake

Our experiments with the purified and reconstituted channels corroborated the mutation-induced proton flux reduction (Fig. 3A). The ratio *R* of the proton flux rate *λ*_D,wt_ per wild-type dimer and the flux rate *λ*_D,D174A_ per mutant dimer was equal to: 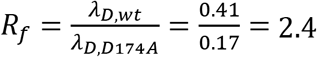 (Fig. 3B).

**Fig. 3.**
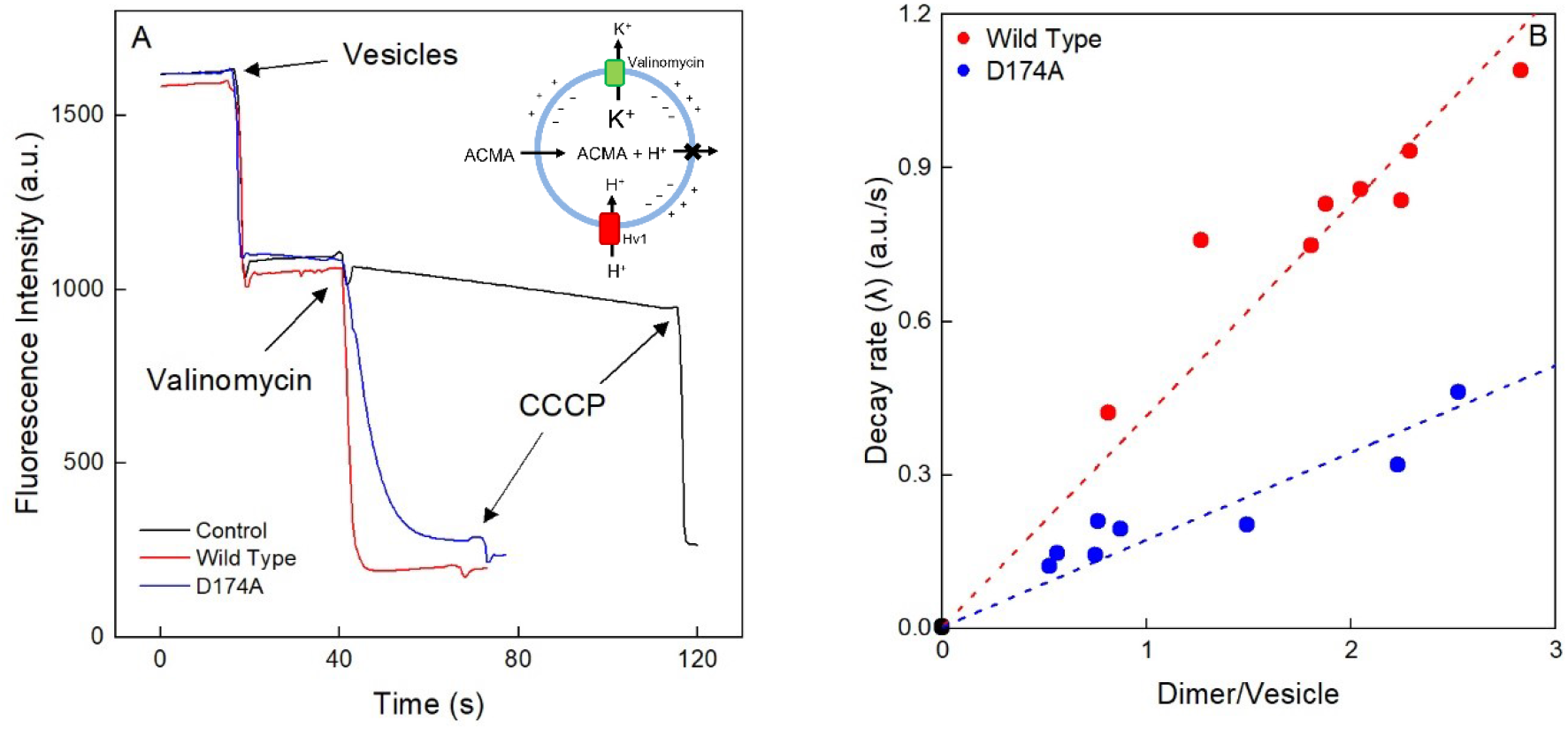
H_V_1-mediated vesicular acidification. A) The addition of vesicles to the dye (ACMA) containing buffer led to a decrease in the fluorescence intensity (*I* in a.u.) due to the (i) dilution and (ii) uptake of the dye into the vesicles. The subsequent addition of valinomycin facilitated K^+^ efflux, which in turn gave rise to membrane potential formation. The latter induced proton uptake via wild-type H_V_1 (red) or the D174A mutant (blue), accompanied by a pronounced fluorescence decrease. The background permeability of the lipid bilayers to protons was much smaller, as indicated by the moderate fluorescence intensity change detected for control vesicles (black). The addition of carbonyl cyanide m-chlorophenyl hydrazone (*CCCP*) revealed the relative size of the vesicle population without H_V_1. The temperature was equal to 23 °C. Inset: Schematic illustration of the fluorescence-based H^+^ flux assay: 150 mM KCl inside the vesicles, 3 mM KCl and 147 mM NaCl outside. B) Proton uptake rate, λ, as a function of protein abundance in the membrane. A linear fit to the data measured for the wild-type H_V_1 (red) and the D174A mutant (blue) yielded the dashed lines. Their slopes indicate a rate λ_D_ = (0.41 ± 0.019) a.u.s^−1^ per wild type dimer and λ_D_ = (0.17 ± 0.013) a.u.s^−1^ per mutant dimer. λ for empty vesicles amounted to λ=0.00196 a.u.s^−1^ (black). λ was determined by fitting a monoexponential function to the *I*=f(time) traces (see panel A). We performed three independent purifications and reconstitutions for each wild type and mutant to obtain λ.

To obtain *λ*_D_, we encapsulated 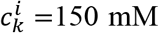 KCl in the H_V_1 containing large unilamellar vesicles (LUVs) and exposed these vesicles to a buffer with a K^+^ concentration 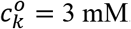. The addition of valinomycin facilitated K^+^ efflux, thereby inducing a membrane potential, *ψ. ψ* constituted the driving force for H^+^ uptake. It can be calculated according to the Goldman equation:

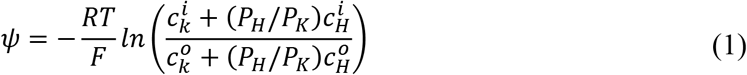

The ratio of the H_V_1 mediated proton permeability *P_H_* to the valinomycin-mediated potassium permeability *P_K_* is always smaller than 0.04. We base our conclusion on the observation that the CCCP-mediated proton permeability represents an upper limit for *P_H_* since CCCP always induces a faster vesicular proton uptake than H_V_1 (Fig. 3). Accordingly, the maximum value of *P_H_/P_K_* can be estimated as the ratio of valinomycin to CCCP conductivities. The respective values are equal to 1.6 10^−3^ Ω^−1^ cm^2^ (*30*) and 4 10^−6^ Ω^−1^ cm^−2^ (*31*). At pH 7.5, we find 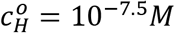, i.e., 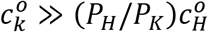. Similarity, 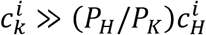 for a broad range of intravesicular pH. With these simplifications, Eq. 1 transforms into the Nernst equation yielding:

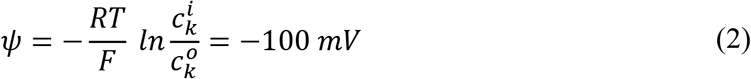

*ψ* of such size may decrease intravesicular pH by nearly two units. Such acidification does not violate 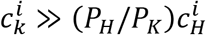 so that *ψ* remains constant throughout the experiment. That is, the vesicle experiments proceed under voltage clamp conditions. The simple explanation is that, due to the small proton concentration and the limited buffer capacity, the K^+^ conductance exceeds H^+^ conductance under all conditions. The conclusion is in line with simulations (*32*), confirming that the membrane potential is driven very near the Nernst potential for K^+^.

To assess the uptake rate, we measured the fluorescence of the weak acid 9-amino-6-chloro-2-methoxyacridine (ACMA). ACMA freely permeates the vesicle membrane (*33*). Yet, when protonated, it becomes trapped in the LUVs, and the fluorescence intensity, *I*, decreases. The dimerization (or oligomerization) of protonated ACMA is most likely responsible (*32*). Fitting a monoexponential function to *I*(t) yielded *λ*. Plotting *λ* as a function of the number of dimers per vesicle (Fig. 3) allowed calculating the transport rate *λ*_D_ per dimer.

### Reconstitution efficiency

The number *N*_C_ of reconstituted H_V_1 dimers per vesicle determines the acidification rate λ, i.e., the time that elapses before reaching the steady state. The final intraluminal pH is independent of *N*_C_. Similarly, CCCP addition in the steady state does not change the intraluminal pH of H_V_1-containing vesicles. But CCCP will affect the intraluminal pH of vesicles deprived of H_V_1 since H^+^ background permeability is too small to allow vesicle acidification within the time allotted for the experiment. Consequently, only H_V_1-free vesicles will acidify upon CCCP addition. That is, CCCP addition allows estimating the fraction of vesicles deprived of H_V_1.

Fluorescence correlation spectroscopy (FCS) revealed *N*_C_ (Fig. 4) (*34*). FCS first counted the number *N*_PL_ of proteoliposomes per confocal volume. Subsequently, a mild detergent (octyl glucoside) dissolved the LUVs and FCS counted the number *N*_MD_ of dimer-containing micelles. After considering the dilution, the ratio *N*_MD_*/N*_PL_ yielded *N*_C_. The subsequent addition of 3 M urea doubled the number of particles, indicating the dissolution of dimers into monomers (Fig. S3).

**Fig. 4.**
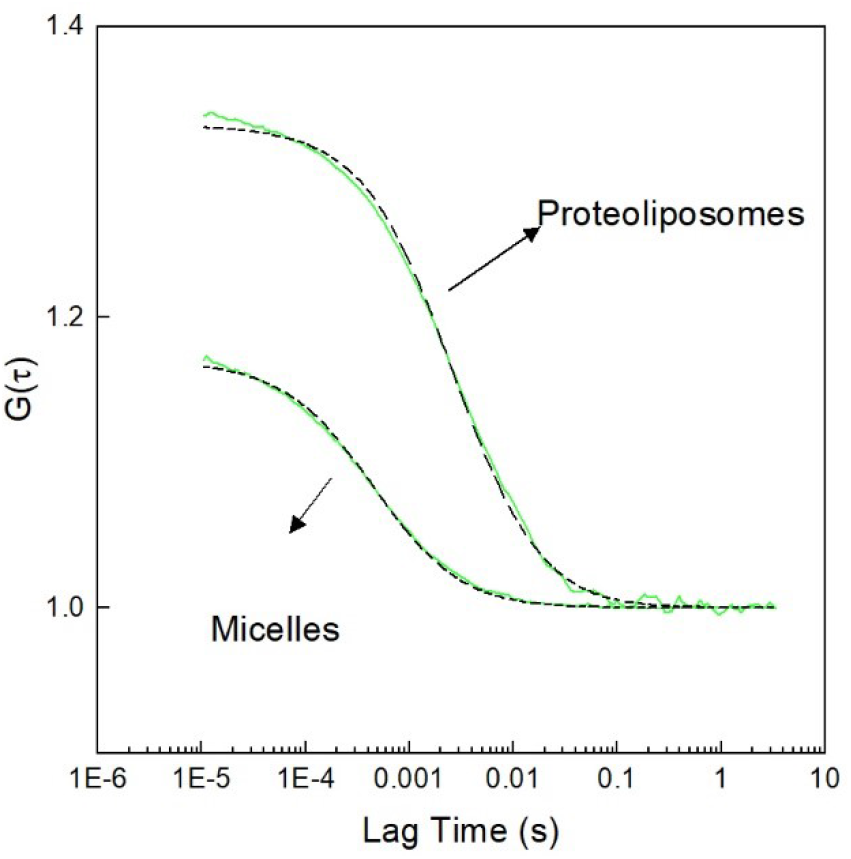
Representative autocorrelation curves *G*(τ) for the determination of reconstitution efficiency. Intensity fluctuations recorded by fluorescence correlation spectroscopy allowed G(τ) calculation. In turn, *G*(τ) served to determine the number *N*_PL_ = 3.0 of proteoliposomes and the number *N*_MD_ = 5.9 of micelles formed per confocal volume after the detergent’s addition. After correcting the dilution during detergent addition, we found the number, *N*_C_, of dimers per vesicle *N*_MD_*/N*_PL_ = 2.25.

### Assessment of the unitary channel conductance

λ_D_ allows assessing the single-channel proton flux. We base our calculation on comparing the known CCCP transport rate because the ACMA assay does not allow precise intravesicular pH determination. The previously reported CCCP mediated conductance, *g_c_*, amounts to about 4×10^−5^ Ω^−1^ cm^2^ for our experimental conditions. Taking into account the transmembrane voltage, *ψ* = −100 mV, we find a current density of:

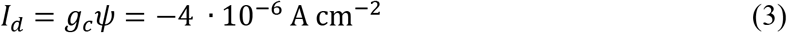

The inward-directed current I_v_ through the membrane of a 100 nm wide vesicle (surface area A_v_) amounts to:

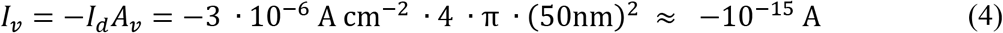

We calculate the number of charges *q* (H^+^) corresponding to *I*_v_ as:

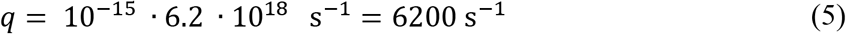

Assuming proportionality between the number of dimers per vesicle and λ (Fig. 3), we estimate that about 4.4 H_V_1 dimers produce the same decay rate, λ = 1.8 a.u. s^−1^, as a CCCP-doped bilayer. Considering the random orientation of the reconstituted channel, only 50 % of them open at the applied voltage. Consequently, we estimate H_V_1’s unitary transport rate, *g_Hv1_*, as:

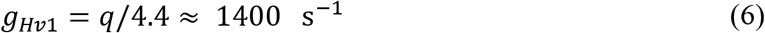

Second, we show that the transport limiting step is H^+^ diffusion to the pore (access resistance) and not transport through the pore. Therefore, we first calculate the maximum current *I*_max_ permitted by diffusion for a constantly open pore (*35*):

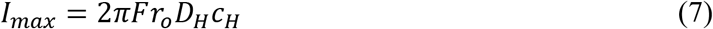

where F, r_0_, D_H_, and c_H_ are Faraday’s constant, the capture radius, the H^+^ diffusion constant, and the H^+^ concentration, respectively. The only unknown parameter is *r*_0_. Taking the gA estimate *r*_0_ = 0.87 Å (*36*), disregarding buffer effects and assuming *D*_H_ = 8.65×10^5^ cm^2^s^−1^, we find:

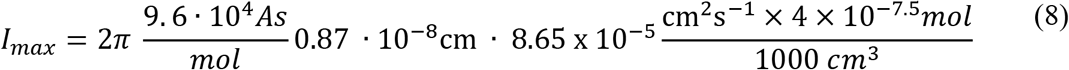

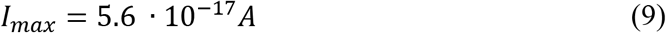

Eq. 8 considers that the approximately 25 % charged lipids in the bilayer induce an increase in surface proton concentration, i.e. it accounts for a surface potential of roughly −40 mV in 153 mM salt. The maximal unitary rate would then be equal to:

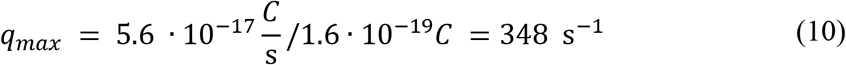

Here we used the *r*_0_ value determined for gA (*36*). Acidic moieties at the entrance of H_V_1 and proton surface migration along the lipid bilayer could serve to increase that value (*37, 38*). The observation *q_max_* ≤ *q* suggests transport limitations by poor proton availability. Calculation of the channel resistance, *R*_ch_ (*35*), confirms the hypothesis:

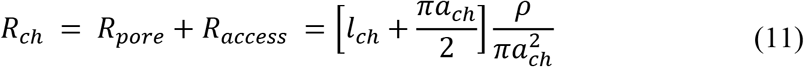

where *R_pore_* is the resistance of the pore proper and *R_access_* is the access resistance. Assuming a channel radius, *a_ch_*, of 0.15 nm, a length, *l_ch_* of 4 nm and solution resistivity (H^+^ as the sole conducted ion at bulk pH of 7.5 and a surface potential of −40 mV), *ρ*, of 2×10^5^ Ω cm, we find *R_ch_* = 4 · 10^13^n. Thus, the resulting current, *I*_ρ_, that we may expect for the vesicular membrane potential of 100 mV is equal to 3 · 10^−15^*A* Accordingly, *I*_ρ_ exceeds *l_max_* by more than one order of magnitude. Consequently, we may safely conclude that H_V_1 conductance is limited by proton availability under our conditions.

### Experiments with the model channel gramicidin

To augment the credibility of the approach exploited for *g_Hv1_* determination, we repeated the procedure for the well know peptide channel gA. We found a roughly fourfold lower *λ*_D_ for gA (Fig. 5).

**Fig. 5.**
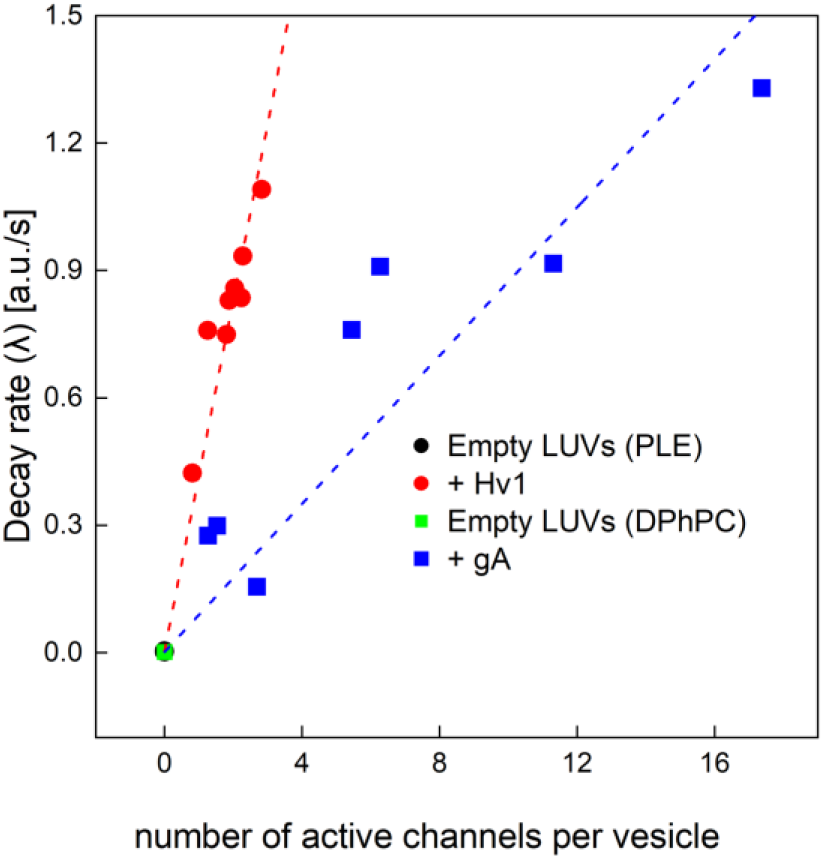
Proton flux through gA channels. We used the same ACMA assay as for H_V_1-containing proteoliposomes. The slope λ_channel_ = 0.09 a.u. s^−1^/channel is fourfold smaller than λ_channel_ = 0.41 a.u. s^−1^/channel for H_V_1 (shown for comparison). The buffer solution contained 150 mM KCl, 5 mM HEPES (pH 7.5), and 0.5 mM EGTA. Substitution of 147 mM KCl for cholinchloride in the external solution drove proton uptake.

The decay rate, λ_D_= 0.09 s^−1^ (Fig. 5) of proton uptake by gA allows calculating gA’s unitary transport rate, *g_gA_* = 310 H^+^ s^−1^ (Eqs. 5–6). Our estimate is reasonably close to single channel conductivities of roughly 90 H^+^ s^−1^ (*39*), which is extrapolated from measurements under very acidic conditions (see also Eq. 7–10 for neutral membranes).

Since the gA channels were unlabeled, we determined their number *n* per vesicle by measuring the increment they introduced to vesicle water permeability, *P*_f_:

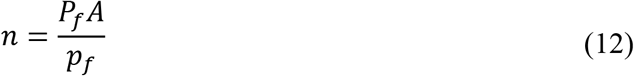

where *p_f_* = 1.6×10^−14^ cm^3^ s^−1^ (*40*) is the unitary channel permeability, and *A* is the surface area of the vesicle (Fig. 6).

**Fig. 6.**
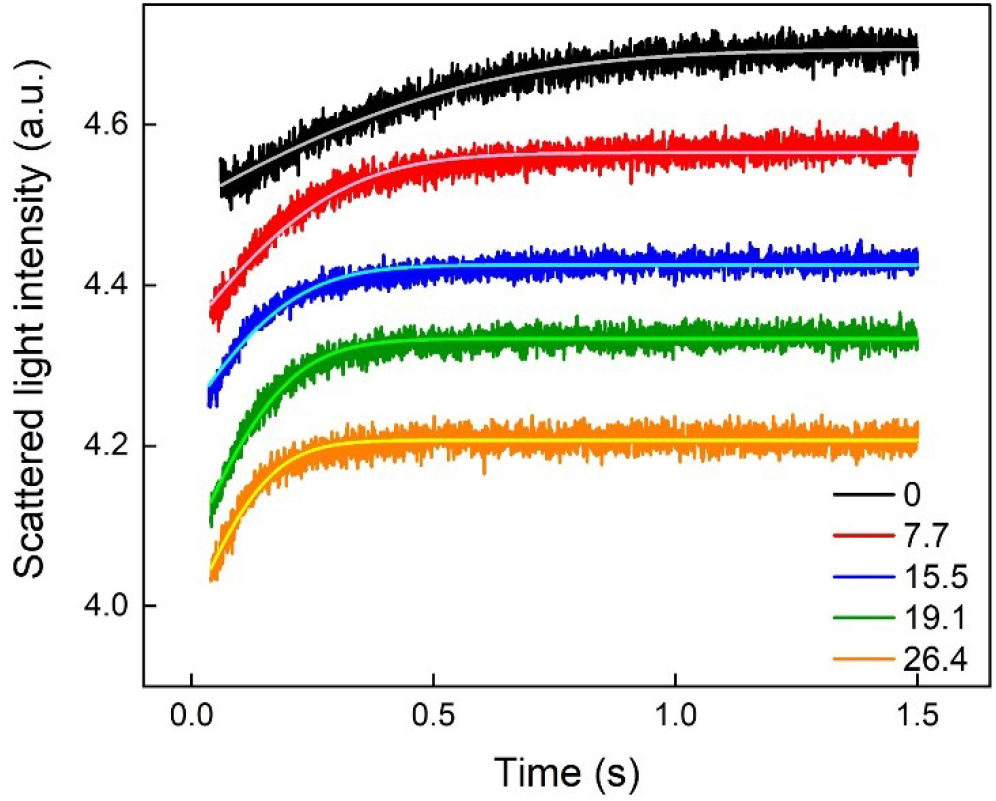
Water flux through gA channels. The bilayer gA concentration decreased from the top (control) to the bottom. The vesicles deflated in response to an osmotic gradient of 150 mmol glucose in a stopped-flow device resulting in an increase of scattered light intensity, *I*_s_. The respective water permeabilities amounted to 2.7, 5.4, 6.1, 7.6, and 8.6 μm s^−1^. The buffer solution contained 150 mM choline chloride, 5 mM HEPES (pH 7.5), and 0.5 mM EGTA.

We calculated *P*_f_ from the osmotically induced volume decrease of gA doped vesicles. The intensity of scattered light *I*_s_ allowed for monitoring the deflation kinetics (compare Materials and Methods).

Since the buffer conditions for the gA water and proton flux masurements differed, we used tryptophan fluorescence in terms of a scale. That is, we first plotted the number of water channels per vesicle as a function of tryptophan fluorescence intensity, *I*_W_, and found *a*_1_ from the linear regression: *I_W_* = *a*_1_ *n* + *b*_1_. Second, we determined the tryptophan fluorescence intensity of the samples used for proton flux measurements. Third, we used *a*_1_ to calculate *n* in these samples. The thus found *n* allowed plotting Fig. 5.

### Water flux measurements with H_V_1 containing vesicles

As before, we measured the intensity of scattered light *I*_s_ to follow the deflation kinetics (Fig. 7). However, neither the proteoliposomes containing the wild type nor the mutant channel showed an increased water permeability, *P*_f_, at room temperature (Fig. S4). This result was independent of the number of reconstituted channels per vesicle. Considering the stark dependence of the activation energy for background water flow across lipid bilayers (*24*), we repeated the experiments at a decreased temperature of 4°C. Thanks to the low background water permeability at 4°C, even tiny contributions of H_V_1 to *P*_f_ should be detectable. Yet, the channels did not contribute to the water flow through the vesicular membrane even though channel water permeability but weakly depends on temperature (*24*). In contrast, both mutant and wild-type channels facilitated H^+^ transport at the decreased temperature (Fig. S5).

**Fig. 7.**
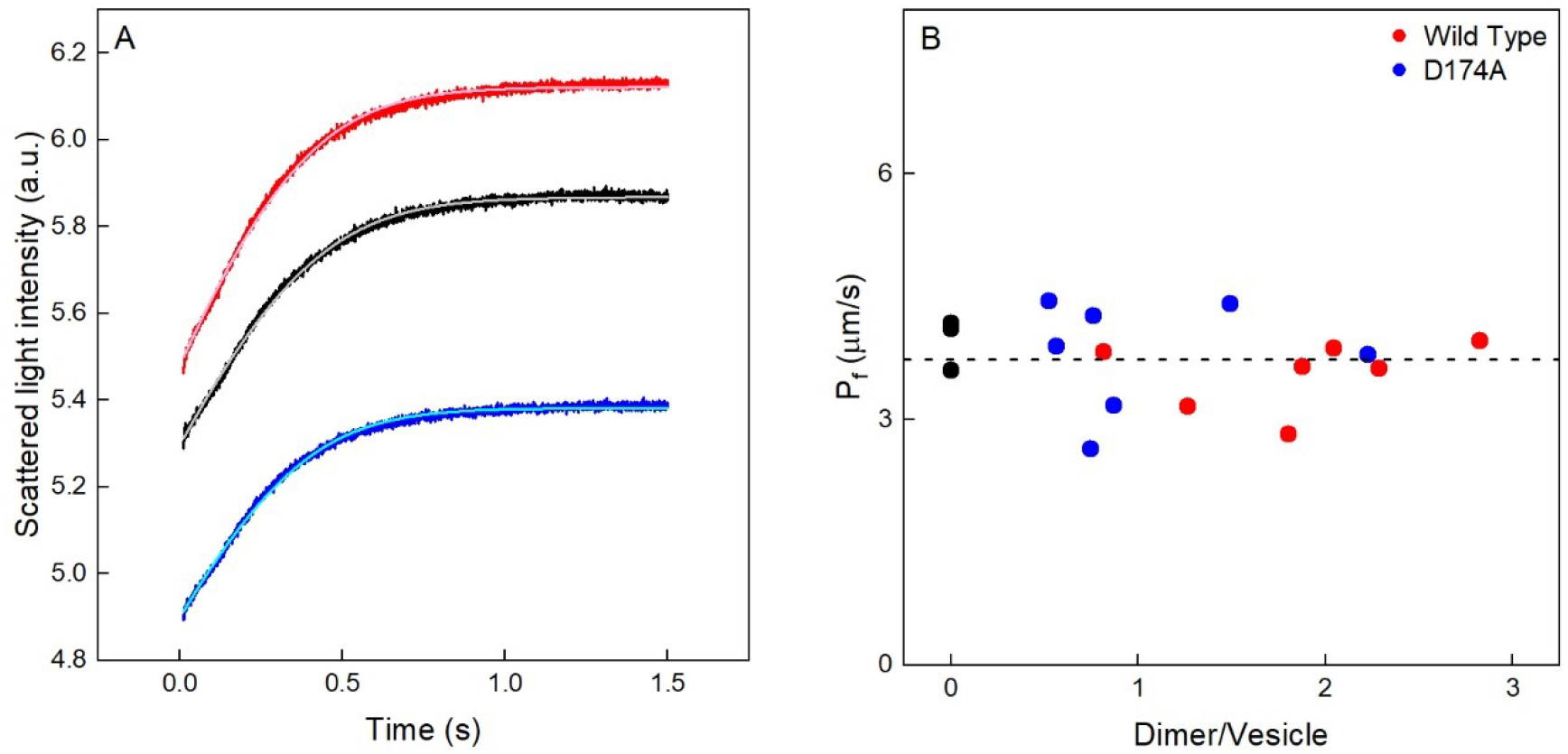
Water permeability of H_V_1 containing LUVs. A) Shrinking of control vesicles (black curve), LUVs containing the D174A mutant (blue), or the wild type (red) induced by the addition of 150 mM sucrose. Otherwise the internal and external solutions were identical: 150 mM KCl, 5 mM HEPES (pH 7.5), and 0.5 mM EGTA. We fitted Equation 20 to the experimental data (grey, light blue, and pink curves) to calculate the vesicular water permeabilities *P*_f_. It amounted to roughly 3.7 μm/s in all three cases. The proteoliposomes contained 0.56 and 0.8 dimers per vesicle. B). Repeating the experiments for higher H_V_1 concentrations also showed no significant *P*_f_ difference between control (filled black circles) and reconstituted LUVs.

## Discussion

Water molecules inside the H_V_1 pore appear to be trapped. They may still serve as proton railways, as demonstrated by the increment in vesicular proton uptake upon channel reconstitution. At the same time, they do not respond to an osmotic driving force, as indicated by the inability of the open channel (D174A) to facilitate water transport across the vesicular membrane (Fig. 8).

**Fig. 8.**
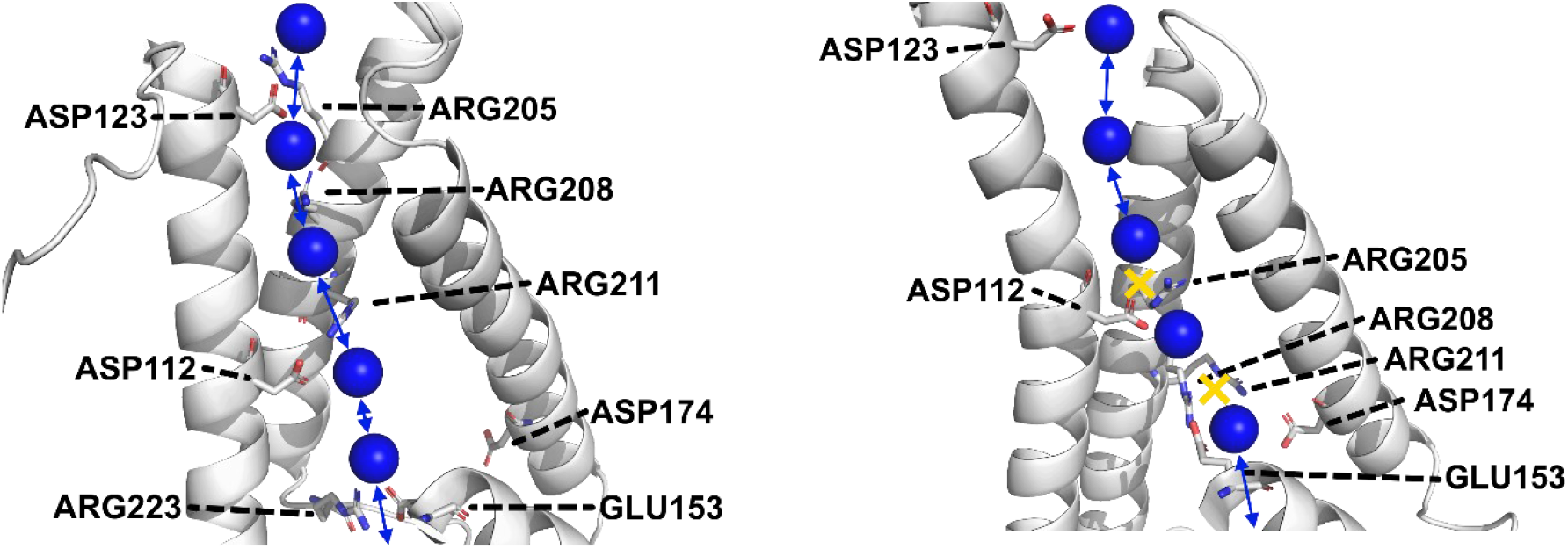
Simplified scheme of proton transport with trapped water molecules inside the channel. It is based on H_V_1 homology models of the open and closed conformations (*13*). A) The open wild-type channel. Salt bridges between the arginines on S4 and acidic residues on S1-S3 may form. In the region of the selectivity filter adjacent to D112, the channel is too narrow to let water molecules pass (see also Fig. S1). Yet, the proton may bypass the electrostatic barrier of the open channel at D112 (*18*), i.e., jump between the two neighboring water molecules. Removal of D174 shifts the voltage sensitivity so that most channels are already open at a transmembrane potential of 0 mv. B) The closed channel. It neither allows water nor proton transport. In its new location, D112 provides an insurmountable electrostatic barrier to proton passage. In contrast to the AlphaFold structure (Fig. 1), the constriction site precluding water movement remains elusive in the homology model (see Fig. S1).

Our result shows that a water wire connecting the solutions on both sides of the membrane does not exist. The contrasting opinion that instead of a channel obstruction hydrogen bonds may immobilize the pore water (*19*) is not convincing. First, the lifetime of a hydrogen bond is in the ps range while H_V_1’s mean open time exceeds 100 ms (*41*). Thus, hydrogen bonds may break more than 10^11^ times during the open state, rendering them unfit for tethering intraluminal water molecules. Second, the effect of hydrogen bonds between water molecules and pore residues is limited to decreased water mobility in narrow channels (*23*). Their number, *N*_H_, allows for predicting *p*_f_ (*26*). Specifically, every H-bond donating or receiving pore-lining residue contributes an average increment ΔΔ*G*^‡^ of 0.1 kcal/mol to the Gibbs free energy of activation Δ*G*^‡^ (*24*). Equation (1) allows the calculation of Δ*G*^‡^:

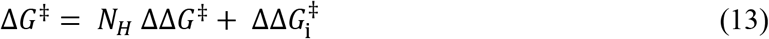

where 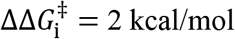 (*24*). Since *N_H_* = 6 (Fig. S1) in the open H_V_1 conformation, Eq. 1 predicts Δ*G*^‡^ = 2.6 kcal/mol. Eq. (2) allows calculating H_V_1’s *p*_f_ from this value (*42*):

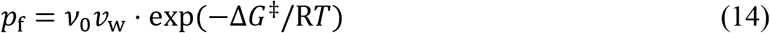

where *v*_w_ = 3 × 10^−23^ cm^3^ is the volume of one water molecule and *v*_0_ is the universal attempt frequency, *v*_0_ = *k*_B_·*T/h* ≈ 6.2 × 10^12^ s^−1^ at room temperature (*k*_B_, is Boltzmann’s and *h* is Planck’s constant). With *p*_f_ = 3 × 10^−12^ cm^3^s^−1^, a single H_V_1 dimer would have contributed 2×*p*_f_/A = 6 × 10^−12^ cm^3^s^−1^/(4 π (50 nm)^2^) = 194 μm s^−1^ to vesicle *P*_f_, where A is the surface area of the vesicle. Upon reconstitution of a single dimer, vesicle *P*_f_ should have increased from 16 μm/s at room temperature (Fig. S4) to 210 μm s^−1^, i.e., by one order of magnitude.

Our gA experiments provide positive control. Since *N*_H_ = 30, the gA dimer has two orders of magnitude smaller *p*_f_ than predicted for the open H_V_1 (*24, 40*). Yet, when gA-containing vesicles are exposed to an osmotic gradient, vesicle deflation is easily discernable (Fig. 6). Thus, in contrast to H_V_1, gA produces the expected *P*_f_ increase. The observation indicates that the H_V_1 pore must be occluded, thereby preventing water passage.

The question arises whether the obstacle in the water pathway is permanent. H_V_1’s titratable residues or steric hindrance from fluctuating sidechains may frequently interrupt otherwise intact water wires. Yet, our calculations (Eq. 7 – 11) show that proton diffusion from the bulk solution to the pore mouth is the transport limiting step. Undoubtedly, transient closure would have caused a detectable pore resistance because part of the protons arriving at the pore mouth could not enter the pore. If the pore was closed longer than one ps, an arriving H^+^ may diffuse out of the capture zone and vanish into the bulk:

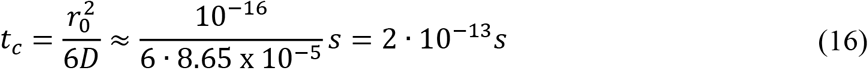

where *t*_c_ denotes the time a proton requires to diffuse a distance equal to the capture radius *r*_0_. Since transient closures would give rise to experimentally undetected pore resistance, they must be ruled out. The observation agrees well with noise experiments, where Lorentzian time constants, albeit smaller than the time constants for H^+^ current activation but larger than 0.1 s were observed (*41*).

The threshold potential to reversal potential relationship is one of the hallmarks of the voltage-gated proton channel H_V_1 (*22, 43*). Here, we characterized the shift of the voltage dependence in the D174A mutant and the resulting negative threshold potential (Fig.2 C+D). Drastically prolonged tail current kinetics might reflect (i) a decreased voltage dependence of the deactivation in the D174A mutant or (ii) a stabilized open state (*14*). Eliminating the negative charge in position D174 2.4-fold decreases the transport rate. The observation supports the hypothesis that D174 is involved in the proton permeation pathway (*12*). The negatively charged entrance point to the pore (Figs. 1B, Fig. 8) may increase *r*_0_ by the same factor it increases λ_D_, i.e., from 0.87 Å measured for gramicidin (*36*) to 2.1 Å. As a result, the area substantially increases from which protons are captured for subsequent channel translocation. Consequently, the diffusion limit increases to 835 H^+^ s^−1^ (Eq 7) – a value within a factor two of our current estimate.

Interestingly, our estimate of 1400 H^+^ s^−1^ is sixfold smaller than a previous approximation of 9400 H^+^ s^−1^. The latter stems from noise measurements of proton currents in inside-out patches of human eosinophils measured between pH 5 and 6.5 (*41*). When extrapolated to physiological pH, the noise analysis yielded a unitary conductance of 15 fS and room temperature, translating into a current of 1.5 fA at a potential of 100 mV. The agreement with our results appears reasonable for the attainable accuracy of noise analysis. Our approach is unlikely to underestimate H_V_1 conductance since it overestimated gramicidin H^+^ conductivity.

In summary, we conclude that our experiments yielded reasonable estimates for single-channel conductivity of reconstituted H_V_1. The value of 1400 H^+^ s^−1^ agrees with the calculated diffusion limit for a ~2 Å capture radius. The high *r*_0_, compared to gA, may be due to the presence of charged amino acids (D174) at the entrance. Neither the closed nor the open state of the proton channel conducts water. The result is in line with models of the closed structure (Alphafold, Fig. 1) and the open structure (homology model, Fig. S1). Both of them show obstructions in the water pathway. The lack of a continuous water wire connecting the solutions on both sides of the membrane indicates that at least one amino acid side chain must be involved in proton conductance.

## Materials and Methods

### Experimental Design

First, we looked for an H_V_1 mutant with a high open probability at zero transmembrane potential. We performed whole-cell recordings of overexpressed human H_V_1 wild type and the D174A mutant.

We assessed H_V_1’s water-conducting ability in a reconstituted system. That is, we purified the overexpressed human H_V_1 channel from insect cells and reconstituted it into large unilamellar vesicles (LUVs). H_V_1 facilitated proton transport as judged by intravesicular pH changes in response to the application of transmembrane potential. Recording the intensity of scattered light allowed quantifying the osmotically induced vesicle deflation.

### Heterologous expression in mammalian cells and electrophysiology

H_V_1-wild type and D174A mutant DNA were synthesized (Eurofins/Genomics, Ebersberg, Germany) and cloned into the pEX-A2 vector using 5’ *Bam*HI and 3’ *Eco*RI restriction sites. Constructs were later sub-cloned into the vector pQBI25-fC3 or pcDNA3.1, using 5’*Bam*HI and 3’ *Eco*RI restriction sites, and GFP laws fused to H_V_1’s N-terminal (*28, 29*).

tSA201 cells (human kidney cell line) were cultivated to 85% confluency in 35 mm culture dishes and transfected with 1.0 μg plasmid DNA using polyethyleneimine (Sigma, St. Louis, MO, USA). After 12 h at 37 °C in 5% CO_2_, cells were trypsinized and replated onto glass coverslips at low density for patch-clamp recordings and measurements done between 2 – 12 hours afterward. We selected cells emitting green fluorescence. The expression level for wild-type and the D174A mutant allowed discarding the contamination of native proton channels. Whole-cell patch-clamp measurements showed no other voltage- and time-dependent conductance at recording conditions.

Whole-cell patch-clamp recordings (HEKA EPC 10, Lambrecht, Germany) were stored on hard discs and analyzed with ORIGIN® (OriginLab, Northampton, MA, USA). Pipettes were pulled from borosilicate capillaries GC 150TF-10 (Harvard Apparatus, Holliston, MA, USA). Tetramethylammonium (TMA^+^) and methanesulfonate (CH_3_SO_3_^−^) constituted the main ions in the whole-cell solutions (pipette and bath) in a concentration range of 90 - 125 mM. We also added 1 mM EGTA, 1 mM Mg^2+^ and 100 mM of a buffer with a pKa close to pH: 2-(N-morpholino)ethanesulfonic acid (MES) at pH 5.5, Bis(2-hydroxyethyl)amino-tris(hydroxymethyl)methane (BIS-TRIS) at pH 6.5 and N,N-bis(2-hydroxyethyl)-2-aminoethanesulfonic acid (BES) at 7.0. Seals were usually > 3 GΩ. Currents are shown without any correction for the leak or liquid junction potentials. We collected the data between 18 to 21 °C. Two methods determined reversal potentials (*V*_rev_): when the threshold of activation (*V*_thres_) was negative to *V*_rev_ it was determined by the zero current, while when *V*_thres_ was positive to *V*_rev_, then *V*_rev_ was determined with the tail current method

### *Purification of* H_V_1

The H_V_1 wild type and mutant D174A were expressed as an N-terminal 10×His-GFP fusion HVCN1 gene in Tni PRO cells (ProTech service, VBCF Protein Technologies Facility, Vienna BioCenter, Vienna, Austria). At 20000 psi, an EmulsiFlex-C5 homogenizer (Avestin, Inc., Ottawa, ON, Canada) disrupted the resuspended cell pellet in a buffer containing 150 mM Tris-HCl (pH 8.0), 500 mM NaCl, and a cocktail of proteinase inhibitors (Roche, Basel, Switzerland). Centrifugation for 10 minutes at 7000 x g (4°C) pelleted the dell debris. 3 M Urea and 5 mM β-mercaptoethanol stripped the membranes within 2 hours at 4°C. Overnight, 1.5% DDM (Anatrace, Inc., Maumee, OH, USA) dissolved the resuspended membrane pellet obtained by ultracentrifugation for 1 hour at 100,000 × g (4°C). 30 minutes of centrifugation at 30000 × g yielded the supernatant subsequently subjected to affinity purification by Ni-NTA resin (Cube Biotech GmbH, Monheim am Rhein, Germany). A washing step with a buffer containing 150 mM Tris-HCl (pH 7.5), 500 mM NaCl, 0.5% DDM, 80 mM imidazole, and proteinase inhibitor cocktail followed 2 h of incubation at 4°C on an over-head shaker (Sunlab Tube roller D-8400, neoLab Migge GmbH, Heidelberg, Germany). The elution buffer contained an additional 0.5 M imidazole. An SDS-PAGE documented H_V_1 purity (Fig. S2). Fluorescence correlation spectroscopy (FCS) determined protein concentration.

### *Reconstitution of* H_V_1

A buffer containing 150 mM KCl, 5 mM HEPES (pH 7.5), EGTA 0.5 mM hydrated a dry lipid film preformed from *E. coli* polar lipid extract (Avanti Polar Lipids, Inc., Alabaster, AL, USA) on the walls of a flask. An ultrasonication bath (Avanti Polar Lipids, Inc., Alabaster, AL, USA) increased the transparency of the 20 mg/ml lipid solution. 40 mM DM primed the lipid mixture for the sequential addition of the H_V_1 detergent mixture. 72 hours of dialysis with tubings containing 15 kDa cutoff pores and subsequent ultracentrifugation (Optima LE-80K, Beckman Coulter Ultracentrifuge, Beckman Coulter Inc., Brea, CA, USA) at 100,000 × g for 2 hours at 4°C removed the detergent. Extrusion through 100 nm pore size filters of the resuspended proteoliposomes yielded uniformly sized large unilamellar vesicles.

### Fluorescence correlation spectroscopy (FCS)

We used FCS to estimate H_V_1 abundance in reconstituted vesicles (*44*). In brief, the FCS unit (Confocor3, Carl Zeiss, Jena, Germany) of a commercial laser scanning microscope (LSM 510) measured the temporal fluctuations of the fluorescence intensity *I* in the focal volume of a C-Apochromat 40x/1.2W objective. The corresponding autocorrelation function *G(τ*) yielded (i) the average number *<N>* of protein-containing particles in the focal volume of radius *r* and elongation *z* of the focus in the direction of the laser beam, and (ii) their diffusion coefficient *D*_v_ (*45*):

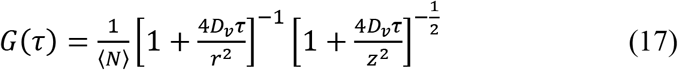

Dissolving the LUVs in mild detergent yielded protein-containing micelles. The ratio of micelles to LUVs indicated the number of dimers per vesicle. Using urea allowed for determining the number of monomers per LUV.

### Proton uptake into LUVs

Transmembrane voltage constituted the driving force for proton uptake into LUVs (Fig. 3A, inset). It resulted from facilitated K^+^ efflux out of the vesicles (*32*). The ensuing vesicular acidification led to the protonation of the dye 9-Amino-6-chloro-2-methoxyacridine (ACMA). Yet the intravesicular concentration of the unprotonated form did not change since it is a weak base that can freely move across the membrane in its neutral form. Intravesicular accumulation of ACMA’s protonated form decreased detectable fluorescence – conceivably due to dye dimerization (*46*).

The extravesicular buffer contained 5 mM HEPES (pH 7.5), 147 mM NaCl, 3 mM KCl, 0.2 mM EGTA and 2 μM ACMA. Sample excitation at 275 nm and measurements of the resulting fluorescence at 475 nm (Fluorescence Spectrophotometer F-2700, Hitachi, Ltd., Chiyoda City, Tokyo, Japan) under vigorous stirring and temperature control (set to 23°C or 4°C) allowed time-lapse experiments. They started by adding 25 μl of a LUV suspension (10 mg lipid per ml) and pipetting 0.5 μl DMSO solution containing 1 mM valinomycin. Additions of 1.5 μM carbonyl cyanide m-chlorophenyl hydrazone (CCCP) equilibrated pH at the end of the experiment.

### Water efflux measurements

Water efflux leads to a decrease in vesicle volume V. The latter can be accessed via measurements of the intensity *I* of scattered light at a wavelength of 480 nm (*23*) according to Equation 3:

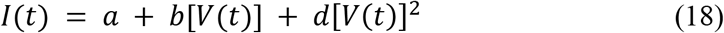

where the parameters *a, b*, and *d* are fitting parameters. We found *a* from Equation (5):

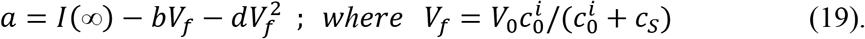

*c*_0_^i^ and *c*_s_ are the initial osmolyte concentrations inside the vesicles and the incremental osmolyte concentration in the external solution, respectively. The rate of the osmotically driven volume decrease determines the vesicular water permeability *P*_f_ (Equation 6):

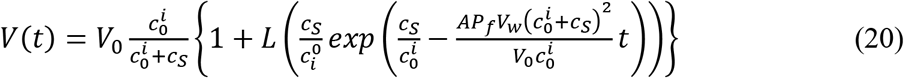

where *V*_w_, *V*_0_, and *L* are the molar volume of water, vesicle volume at time zero, and the Lambert function: *L*(*x*)*e*^*L*(*x*)^ = *x*, respectively (*23*).

The system of Equations (18 - 20) was globally fitted to the whole set of shrinking curves obtained in a stopped-flow device (SFM 3000, Bio-Logic, Seyssinet-Pariset, France) from a particular reconstitution sample to extract the fitting parameters *b*, *d*, and *P*_f_.

## Author contributions

Conceptualization: BM, PP.

Methodology: CS, CH

Investigation: DB, SB, GC.

Visualization: SB, GC, SK, NGM.

Supervision: BM, PP.

Writing—original draft: PP

Writing—review & editing: DB, SB, GC, CH, CS, SK, NGM, BM, PP

## Data and materials availability

All data are available in the main text or the supplementary materials.

