## Supplemental Material for "Trapped pore waters in the open proton channel H_V_1"

The Supplement contains five figures: Fig. S1 contains structural models of the open and the closed states. Fig. S2 shows SDS and fluorescence gels. Fig. S3 displays representative records from fluorescence correlation spectroscopy that show how we determined the number of channels per vesicle. Fig. S4 shows vesicular water efflux at 4°C and 23°C. It demonstrates the inability of both the closed wild-type channel and the open D174A mutant to transport water. Fig. S5 Indicates that the water-impermeable channels facilitate proton transport.

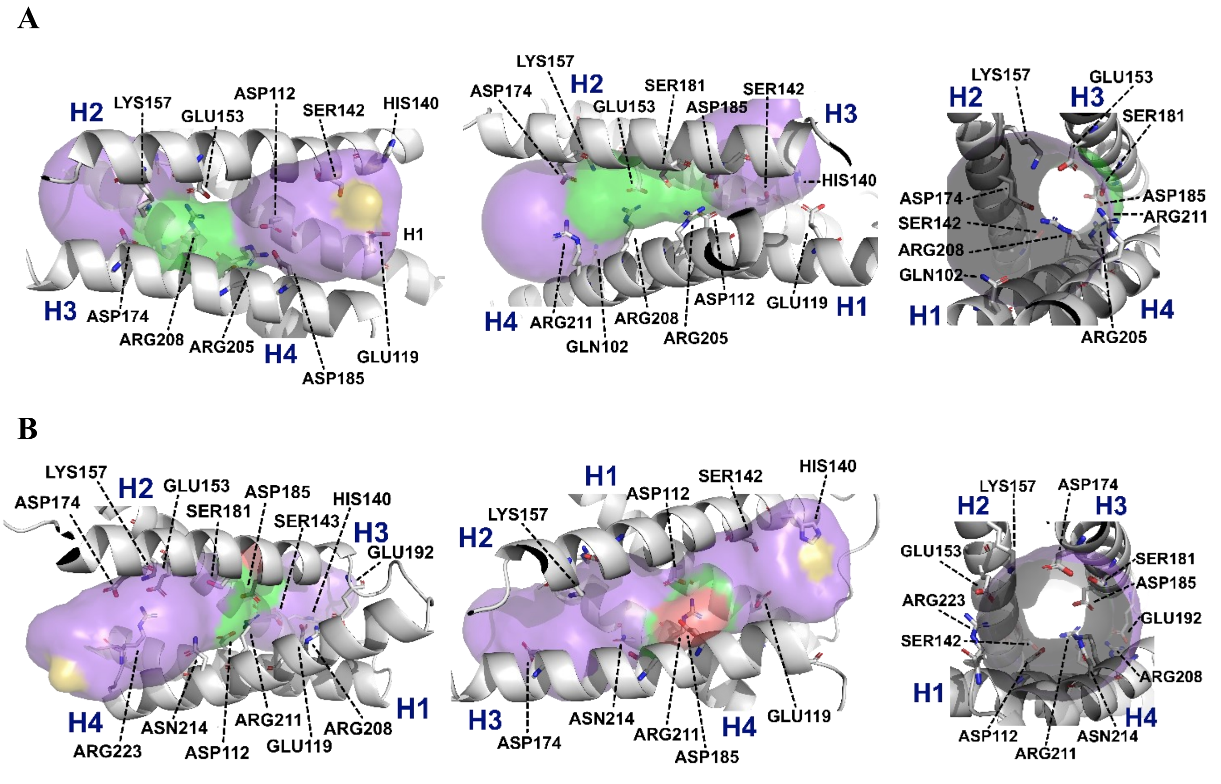


**Fig. S1**. Structural models of the Hv1 channel in the closed state (A) and the open state (B) obtained by a 10-µs-timescale atomistic molecular dynamics simulation in an explicit membrane environment

(*1*). View at the whole channel – cyto: left, peri: right (Left) and turned for 90° around the central channel axis (Middle). View along the channel - cyto > peri (Right). We used the programs PyMol and HOLE to prepare the figure and visualize the channel volume, respectively. The single file region of the channels is colored in green. Red regions mark pore sections too narrow to let a water molecule pass. Pore regions wide enough to let water molecules overtake each other are labeled in violet. The following 10 amino acid dip into the single file region or border with it: 102, 112, 157, 174, 153, 181, 185, 205, 208, 211 in the closed state. In the open state the single file region is shorter. It consists of only six amino acids: 112, 143, 181, 185, 211, and 214. The HOLE program suggests that the closed structure is permeable to water, whereas the open state shows an obstruction at ASP 185 that prevents water from passing through.

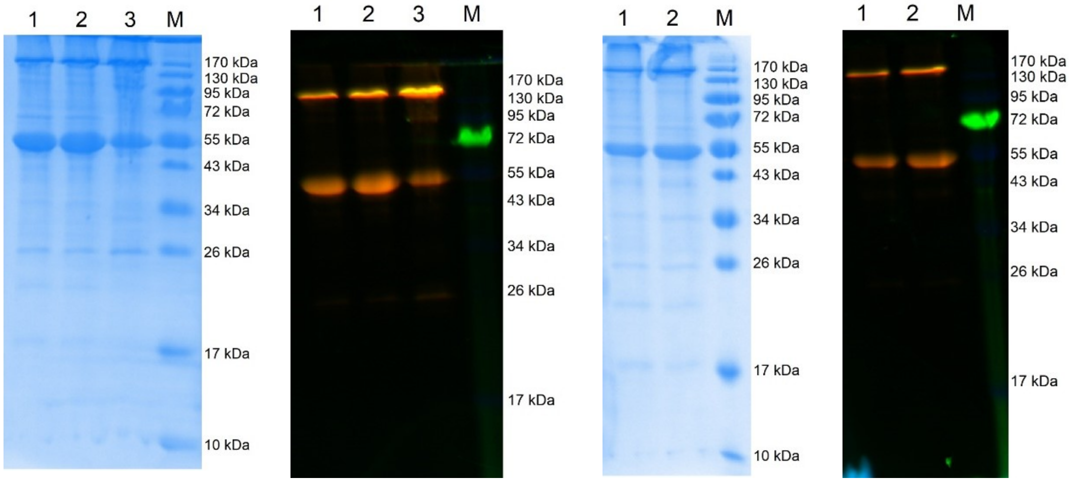


**Fig. S2. Purification of Hv1.** Representative SDS-PAGE runs with purified wild-type Hv1 (A/B) and Hv1 mutant D174A (C/D). The strong bands observable at the height of the marker band at 55 kDa show Hv1 with GFP fused to its N-terminus. A second band can be seen in the range of 130 to 170 kDa, representing Hv1 dimers, likely running higher than expected due to a retained non-globular shape (panel A – wt, and C – D174A). In both cases, presence of GFP in the respective monomer and dimer bands is confirmed via fluorescence detection at 488 nm (panels B and D).

22

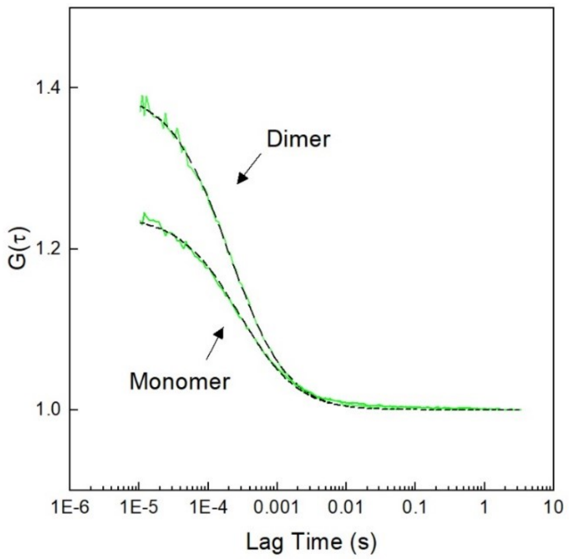


**Fig. S3. Dissolution of HV1 wild-type dimers into monomers.** Representative autocorrelation curves $G(\tau)$ showing HV1 dimers in micelles formed by the addition of mild detergent to proteoliposomes. Subsequent addition of 3 M urea yielded monomers. $G(\tau)$ indicates the number of monomers $N_{MM}$ per confocal volume to be equal to 4.5 and the number $N_{MD}$ of dimers to be equal to 2.6. Correcting for the dilution by urea, yields the number of monomers per dimer $N_{MD}/N_{MM} = 2.1$

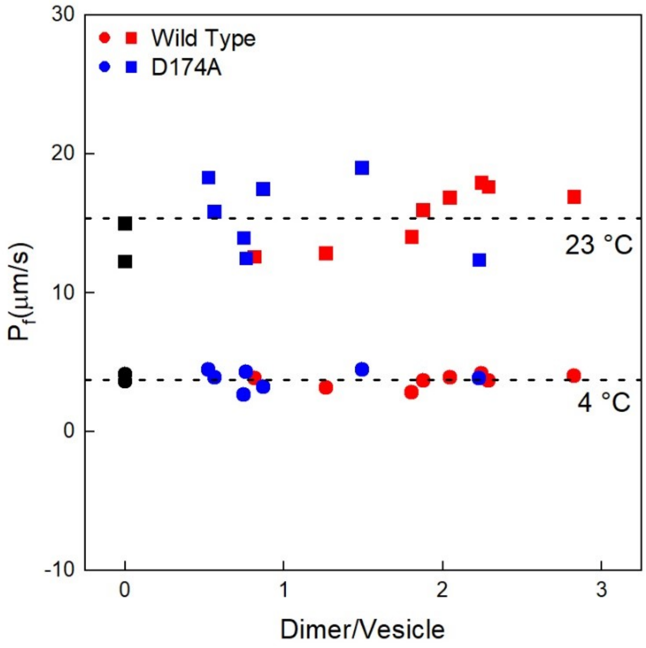


**Fig. S4. Hv1 does not facilitate water transport.** We repeated the experiment shown in Fig. 5 at 23°C (squares). The results shown in Fig. 5 are redrawn (circles). Lipid vesicles without the protein channel (black) have the same water permeability, $P_f$, as those containing the wild-type protein (red) and the mutant D174A open at 0 mV.

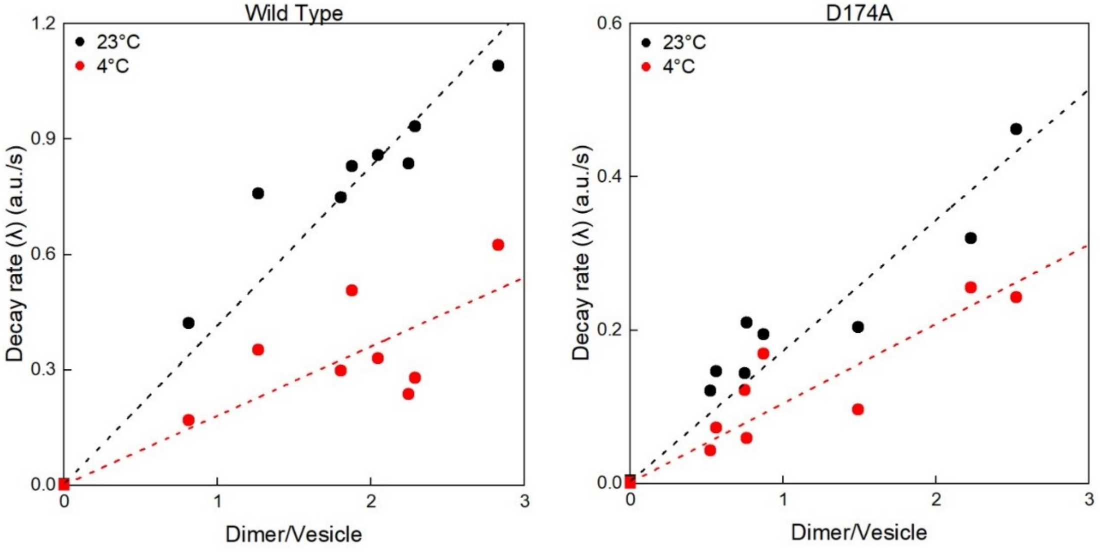


**Fig. S5. Hv1 facilitates proton transport at a decreased temperature.** We repeated the experiments shown in Fig. 3 at a reduced temperature of 4°C. The comparison to the results at room temperature (redrawn from Fig. 3) shows a roughly twofold drop of activity for both wild-type and mutant channels upon a temperature decrease of 19 °C.
